# Propagation of viral genomes by replicating ammonia-oxidising archaea during soil nitrification

**DOI:** 10.1101/2022.08.13.503859

**Authors:** Sungeun Lee, Ella T. Sieradzki, Graeme W. Nicol, Christina Hazard

## Abstract

Ammonia-oxidising archaea (AOA) are a ubiquitous component of microbial communities and dominate the first stage of nitrification in some soils. While we are beginning to understand soil virus dynamics, we have no knowledge of the composition or activity of those infecting nitrifiers or their potential to influence processes. This study aimed to characterise viruses having infected autotrophic AOA in two nitrifying soils of contrasting pH by following transfer of assimilated CO_2_-derived ^13^C from host to virus via DNA stable-isotope probing and metagenomic analysis. Incorporation of ^13^C into low GC mol% AOA and virus genomes increased DNA buoyant density in CsCl gradients but resulted in co-migration with dominant non-enriched high GC mol% genomes, reducing sequencing depth and contig assembly. We therefore developed a hybrid approach where AOA and virus genomes were assembled from low buoyant density DNA with subsequent mapping of ^13^C isotopically enriched high buoyant density DNA reads to identify activity of AOA. Metagenome-assembled genomes were different between the two soils and represented a broad diversity of active populations. Sixty-four AOA-infecting viral operational taxonomic units (vOTUs) were identified with no clear relatedness to previously characterised prokaryote viruses. These vOTUs were also distinct between soils, with 42% enriched in ^13^C derived from hosts. The majority were predicted as capable of lysogeny and auxiliary metabolic genes included an AOA-specific multicopper oxidase suggesting infection may augment copper uptake essential for central metabolic functioning. These findings indicate virus infection of AOA may be a frequent process during nitrification with potential to influence host physiology and activity.

## Main text

Microbially mediated oxidation of ammonia to nitrate during nitrification is a central component of the global nitrogen (N) cycle. It is also responsible for major losses of applied fertiliser N in soil, generating atmospheric pollution via direct and indirect production of nitrous oxide (N_2_O) as well as nitrate (NO_3_-) pollution of groundwater [1]. Autotrophic ammonia-oxidising archaea (AOA) of the class *Nitrososphaeria* are a ubiquitous component of soil microbial communities and often dominate ammonia oxidation and nitrification-associated N2O emissions when ammonia is supplied at low rates via organic matter mineralisation [2], slow-release fertilisers [3] or in acidic soils [4]. Integrated temperate viruses (proviruses) and other virus-associated protein-encoding genes are found in most AOA genomes suggesting frequent interaction (see Supplementary Text). While viruses infecting marine AOA have been characterised through metagenomic approaches [5] and cultivation [6], those infecting soil AOA or other nitrifier groups are currently uncharacterised.

Virus infection can influence biogeochemical cycling by augmenting host activity or causing cell mortality and subsequent release of nutrients [7]. Recent advances have demonstrated that soil virus communities are dynamic in a wide range of soils [e.g. 8, 9] and augmenting virus loads modulate C and N fluxes [10, 11]. Nevertheless, identifying active interactions with specific populations or functional groups in soil remains challenging due to structural complexity and the vast diversity of hosts and viruses. Recent use of stable-isotope approaches has investigated whole community host-virus dynamics [12, 13] or interactions between individual host-virus populations specific to a functional process and substrate [14]. The aim of this study was to utilise the latter approach with ^13^CO_2_-based DNA-SIP to focus on nitrification-associated interactions and to test the hypothesis that viruses are a dynamic component of soil AOA activity.

### Hybrid analysis of GC mol% fractionation and ^13^C-DNA-SIP for identifying active AOA hosts and viruses

Microcosms were established (see Supplementary Text) with two sandy loam soils of contrasting pH (4.5 or 7.5), which were characterised previously as containing distinct AOA communities [15]. As ammonia from hydrolysed urea is used by AOA in these and other soils [1, 3], nitrification was fuelled by addition of urea (two pulses of 100 μg N g^-1^ soil (dry weight)) to stimulate AOA growth and potential enrichment of AOA-infecting viruses. Microcosms were incubated at 25°C for 30 days (Supplementary Figure 1) with a 5% ^12^C- or ^13^C-CO_2_ headspace to target viruses of autotrophs via transfer of assimilated carbon. Extracted DNA was subjected to isopycnic centrifugation in CsCl gradients, in which DNA migrates as a function of GC mol% and isotopic enrichment, before fractionation and analysis of genomic DNA of different buoyant densities [14]. Quantification of AOA genomic DNA distribution across CsCl gradients demonstrated growth of AOA populations (Figure 1A; Supplementary Figure 2) and genomic DNA in high buoyant density (HBD) fractions (>1.719 g ml^-1^) was then pooled for each replicate microcosm and metagenomes sequenced (Supplementary Table 1). While ^13^C-incorporation clearly separated DNA from growing vs non-growing AOA populations, ^13^C-enriched AOA genomes (average contig GC mol% content of 38%) co-migrated within the CsCl gradient with more abundant community ^12^C genomes with higher GC mol% (>57%), reducing AOA host and virus sequencing depth, contig assembly and binning. Consequently, only one medium-quality AOA metagenome-assembled genome (MAG) was recovered from HBD DNA and no AOA-associated virus contigs were identified amongst 290 viral operational taxonomic units (vOTUs) ≥10 kb (Figure 1B).

**Figure 1.**
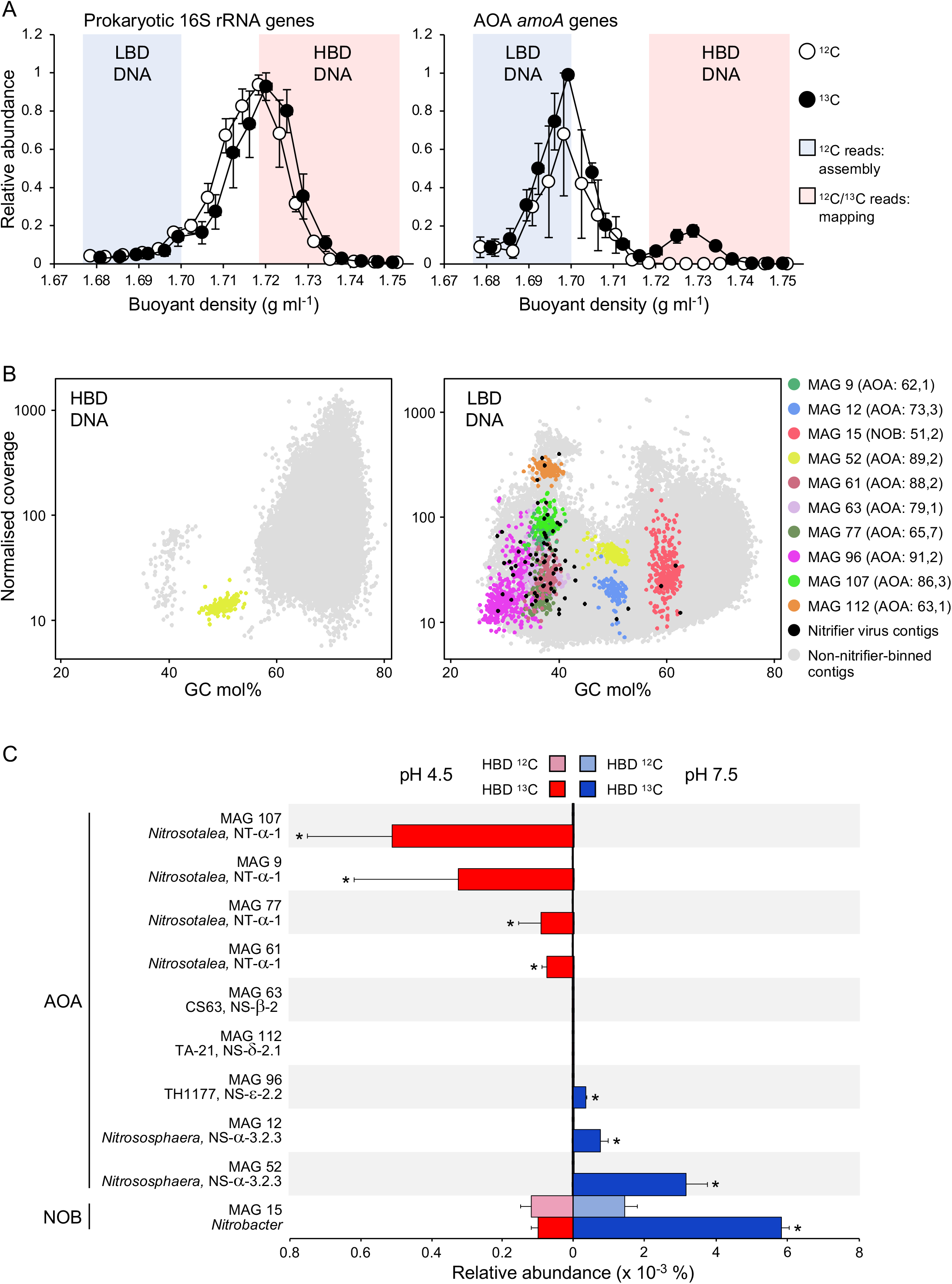
Contigs and MAGs from low GC mol% DNA and *in situ* growth of AOA in pH 4.5 and 7.5 soils determined from hybrid analysis of ^12^C- and ^13^C-enriched DNA. **A** Distribution of total prokaryotic and AOA genomes in CsCl gradients determined from the relative abundance of 16S rRNA genes and *amoA* genes, respectively. Example profiles are shown for pH 4.5 soil only (see Supplementary Figure 2 for pH 7.5 profiles). Vertical error bars are the standard error of the mean relative abundance and horizontal bars (mostly smaller than the symbol size) the standard error of the mean buoyant density of individual fractions from three independent CsCl gradients, each representing an individual microcosm. Genomic DNA in fractions highlighted in blue or pink areas were pooled for each replicate microcosm for metagenomic sequencing. **B** Distribution of assembled contigs (≥5 kb) from metagenomic libraries prepared from LBD or HBD DNA in sequence coverage vs. GC mol% plots. Contigs binned into ten medium- and high-quality nitrifier MAGs and predicted viruses of nitrifiers (≥10 kb) are highlighted, with number in parenthesis beside each MAG identifier giving estimated completeness and contamination (%). **C** Mean and standard error of the relative proportion of metagenomic reads from ^12^C- and ^13^C-enriched HBD DNA mapped onto contigs of nine AOA and one *Nitrobacter* MAGs. A significantly greater relative proportion of reads in ^13^C-derived libraries are indicated with * (*p* <0.05, two-sample Student’s t-test or Welsch’s t-test when variances were not homogenous). GTDB genus [16] and *amoA* gene lineage-affiliation [17] are given.

To facilitate analysis of low GC mol% genomes while also identifying activity, we developed a hybrid approach combining metagenomic analysis of low buoyant density (LBD) ^12^C DNA with read-mapping of sequenced HBD DNA (Figure 1; Supplementary Figure 2). The AOA and virus genomes were assembled from metagenomic libraries of ^12^C LBD DNA only (<1.699 g ml^-1^) representing both dormant and active populations. Individual sequence reads from HBD DNA libraries from both ^12^C- and ^13^C-incubations were then mapped to identify activity with a significantly greater relative abundance in ^13^C-incubated samples demonstrating isotope incorporation. For taxonomically defined contigs of hosts or host-linked viruses, in comparison to HBD, assemblies from LBD libraries increased the relative proportion of contigs from the AOA-containing *Thermoproteota* phylum (GTDB taxonomy; *Nitrososphaerota* in ICNP taxonomy and *Thaumarchaeota* in NCBI taxonomy) and *Thermoproteota-infecting* virus contigs from 0.3 to 2.8% and 0 to 22.3%, respectively (Supplementary Figure 3). In LBD metagenomes, 123 medium- and high-quality MAGs from both soils were assigned taxonomy (Supplementary Table 2) including nine AOA MAGs representing six genera (GTDB classification [16]) within the *Nitrososphaerales* and *Nitrosopumilales* orders and one nitrite-oxidising bacterium (NOB) *Nitrobacter* MAG (MAG 15) (Figure 1B, Supplementary Figure 4), contributing to the oxidation of nitrite (NO_2_-) to NO_3_- during the second step of canonical nitrification. Mapping of HBD sequence reads revealed that eight of the ten MAGs were derived from active autotrophic populations (Figure 1C). Consistent with previous studies characterising the distribution of soil AOA from different taxa [17], this included four *Nitrosotalea* genus populations (MAGs 9, 61, 77 and 107) from pH 4.5 soil and three *Nitrososphaera* genus populations (MAGs 12, 52 and 96) from pH 7.5 soil. No evidence of autotrophic growth was observed for populations represented by a further two MAGs within the *Nitrososphaerales*. Both are from genera with no cultivated representatives with MAG 112 from genus TA-21 (*amoA*-gene lineage NS-ß-2 [17]) and MAG 63 (*amoA*-gene lineage NS-δ-2.1) representing a novel genus (designated here as CS63). The latter belongs to the *amoA* gene-defined NS-δ lineage that while widely found in soil, is rarely implicated in contributing to ammonia oxidation. Although these two MAGs may simply be from AOA populations not active under the incubation conditions used, they are also consistent with microorganisms possessing physiologies distinct from cultivated AOA.

### AOA viruses

After predicting contigs of virus origin using established bioinformatic tools, those from AOA-infecting viruses were identified with a custom database of hallmark genes from AOA proviruses and analysis of other shared homologues (see Supplementary Text). Specifically, proviruses were identified in 184 (87%) of 212 AOA genome sequences downloaded from GTDB. Capsid, terminase, portal and integrase protein sequences from these proviruses were subsequently used as search queries. While previous analysis of 12 total soil metagenome or virome libraries from the same soil samples did not enable identification of any AOA-associated virus contigs (average of 152 million quality filtered reads and 1,947 contigs ≥10 kb per library) [14], this targeted approach using LBD DNA from nitrifying microcosms yielded 64 contigs predicted as derived from AOA-infecting viruses and each representing a unique vOTU. In addition, viruses infecting other nitrifier populations were identified with five and three vOTUs derived from viruses infecting ammonia-oxidising bacteria (AOB) and NOB, respectively. Virus and host genome GC mol% correlate [18] and AOA-infecting virus contigs possessed a similar average content of 36.3% which was considerably lower than averages of 46.6% and 67.0% of all viral contigs recovered from LBD and HBD metagenomes, respectively. As with their hosts, AOA virus communities were distinct between the two pH soils (Figure 2A) as also observed previously for total virus communities [19] with 29 (45%) and 34 (53%) vOTUs significantly more relatively abundant in pH 4.5 and 7.5 LBD libraries compared to the other soil, respectively. Comparison of the relative abundance of HDB sequence reads from both ^12^C and ^13^C libraries revealed that 27 contigs from both soils (42%) were enriched in ^13^C and thus confirming infection during incubation and transfer of C from AOA hosts to viruses.

**Figure 2.**
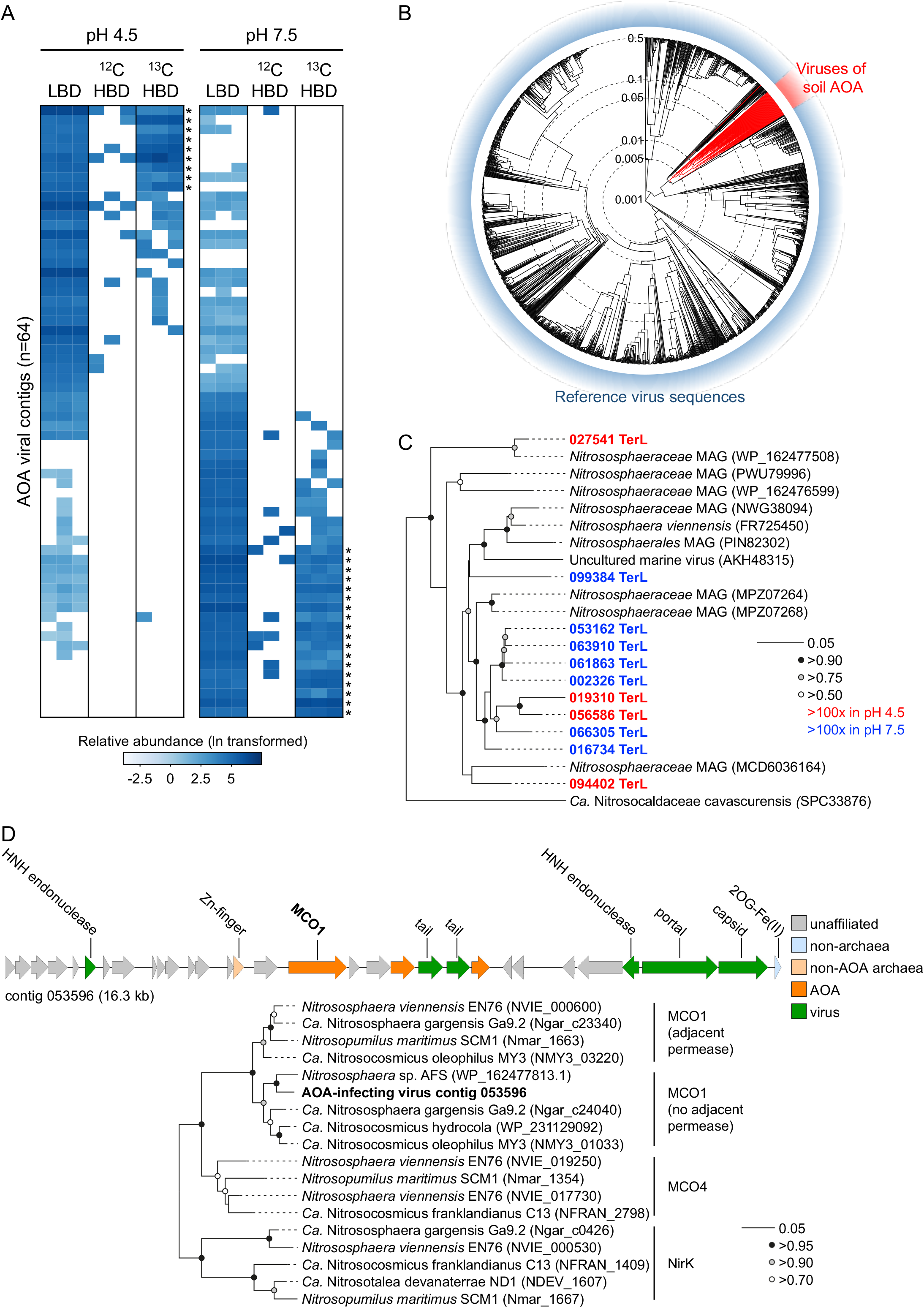
Identification and determination of activity of viruses infecting AOA in pH 4.5 and 7.5 soils using a hybrid analysis of ^12^C- and ^13^C-enriched DNA. **A** Heatmap showing the relative abundance of 64 vOTUs identified in LBD DNA libraries from pH 4.5 and 7.5 soil (≥1× coverage, ≥75% contig breadth). To identify activity, reads from ^12^C- and ^13^C-enriched HBD DNA were mapped onto LBD-derived contigs with a significantly greater relative proportion of reads in ^13^C-derived libraries indicated with * (*p* <0.05, two-sample Student’s t-test or Welsch’s t-test when variances were not homogenous). **B** Proteomic tree showing genome-wide sequence similarities between AOA viral contigs and 1,981 curated virus genomes. Values at dotted circles represent a distance metric based on normalized tBLASTx scores plotted on a log scale. **C** Maximum likelihood phylogenetic analysis of derived large sub-unit terminase (TerL) protein sequences (336 unambiguously aligned positions, LG substitution model) identified in this and other metagenomic studies or in proviruses of cultivated AOA. NCBI accession numbers are given in parenthesis. Circles at nodes represent percentage bootstrap support from 1000 replicates, scale bar denotes an estimated 0.05 changes per position and colour-coding describes the comparative relative abundance in each soil. **D** Genetic map of AOA-infecting virus-derived contig 053596 containing a type 1 multicopper oxidase (MCO1) gene. Maximum likelihood phylogenetic tree describes placement of MCO1 within the AOA MCO/NirK family of cultivated AOA following designations of Kerou *et al.* [22] (206 unambiguously aligned positions, LG substitution model, invariant and gamma distributed sites). NCBI accession numbers are given in parenthesis. Circles at nodes represent percentage bootstrap support from 1000 replicates and the scale bar denotes an estimated 0.05 changes per position.

Soil AOA viruses were distinct from characterised archaeal and bacterial viruses. Gene-sharing network analysis using vConTACT v2.0 [20] did not identify any associations with reference sequences or recently described viruses of marine AOA [6] and genome-wide sequence comparisons [21] placed 60 of the 64 soil AOA viral contigs within a unique proteomic grouping (Figure 2B). The majority were predicted as capable of lysogeny with 55 encoding an integrase. While it can be difficult to determine virulent versus temperate capability from partial genomes, only 6.8 and 8.9% of all virus contigs from LBD and HBD metagenomes possessed a predicted integrase. Although suggesting a higher proportion of AOA viruses may be temperate, it should be noted that this could be a consequence of using AOA provirus-associated genes to identify viral contigs, biasing towards identification of those from temperate viruses. These results also highlight that while DNA-SIP of total metagenomes enables identification of ^13^C incorporation into virus genomes, it is potentially limited in facilitating confident bioinformatic prediction of free temperate virus particles versus proviruses present from a previous infection of a currently active host. While the integration status of a temperate virus can be predicted bioinformatically, using targeted “viromes” metagenomes of filtered virus particles will provide greater confidence of identifying genomes from free virus particles.

Eleven contigs contained large terminase (TerL) sub-unit gene sequences and were similar to those recovered from other predicted AOA virus or provirus genomes only (e-value <10^-5^) (Figure 2C). In addition to viral hallmark genes encoding portal, endonuclease, tail and capsid proteins, contig 053596 contained a type 1 multicopper oxidase (MCO1) homologue found exclusively in AOA genomes [22] (Figure 2D). Copper (Cu) is essential for ammonia oxidation and electron transport in AOA and MCO1 is upregulated under Cu limitation and potentially involved in Cu sequestration from the environment [23]. This virus-encoded MCO1 homologue therefore represents an auxiliary metabolic gene that may contribute to a core process in AOA physiology.

### Summary

These results from two soils indicate that virus infection of AOA during nitrification is likely a common and dynamic process, with the potential to influence nitrification rates in soil. Although viruses were only identified from contigs representing partial viral genomes, analysis of individual marker genes and genome-wide similarity comparisons revealed that those infecting AOA are highly divergent from known viruses. While one *Nitrobacter* MAG was obtained and both AOB and NOB viruses were also predicted, the hybrid approach developed in this study targeted low GC mol% DNA and was specifically tailored for the analysis of AOA populations and their viruses. A broader analysis of all viruses infecting nitrifying prokaryotic populations would therefore likely require a more comprehensive sequencing effort of all fractions throughout the CsCl gradient using quantitative SIP metagenomics [24] or alternative approaches, such as virus-targeted metagenomic viromes.

## Data availability

Metagenome sequence data are deposited in NCBI’s GenBank under BioProject accession nos. PRJNA621418-PRJNA621429 (^13^C data) and PRJNA868779 (^12^C data) with MAGs available under accession numbers SAMN30436125-SAMN30436246.

## Acknowledgements

This work was funded by an AXA Research Chair in Microbial Ecology awarded to GWN, Agence Nationale de la Recherche grant ‘CONSERVE’ (ANR-22-CE02-0006-01) awarded to GWN and CH, and an EU Horizon Europe MSCA Postdoctoral Fellowship ‘DIVOBIS’ awarded to ETS. Sequencing data generated under JGI Community Science Program Proposal No. 503702 awarded to GWN and CH was conducted by the US DOE JGI, a DOE Office of Science User Facility, and supported by the Office of Science of the US DOE under Contract No. DE-AC02-05CH11231. The authors would like to thank Dr. Willm Martens-Habbena (University of Florida) for continued discussions on soil AOA ecophysiology and taxonomy, Dr. Melina Kerou (University of Vienna) for guidance on AOA MCO diversity, Dr. Gary Trubl for comments on the manuscript and Dr. Robin Walker (SRUC, Aberdeen) for access to the much-lamented Craibstone Estate Woodlands Field long term experiment.

## Author contributions

SL, GWN and CH conceived and designed the experiment with additional input from ETS. SL performed all laboratory work and analysed the data with input from GWN in phylogenetic analyses. SL, GWN and CH drafted the manuscript text and figures with all authors contributing to the final paper.

