## Supplementary material for "Propagation of viral genomes by replicating ammonia-oxidising archaea during soil nitrification": Text, tables and figures

**Supplementary text**

**Materials and methods**

*Soil microcosms, DNA extraction and stable-isotope probing*

Soil microcosms were established in 144 ml serum vial bottles with 10 g soil (dry weight equivalent). Microcosms were amended with urea solution (100 µg urea-N $g^{-1}$ soil (dry weight)) at day 0 and 15 with the water content being 30% and 32% (w/w) after amendment, respectively. Headspace gas was amended with 5% (v/v) $^{12}C$-$CO_2$ (Air Liquide) or $^{13}C$-$CO_2$ (Sigma-Aldrich) (99% atom enriched) with microcosms opened every 3-4 days to maintain aerobic conditions before re-establishing $CO_2$ concentrations. Microcosms were destructively sampled at day 0, 15 and 30 in triplicate for each treatment and soil stored at -20°C. Ammonium, nitrite and nitrate concentrations were determined immediately after sampling using standard colorimetric assays [1]. Total genomic DNA was extracted from 0.5 g soil using a CTAB-buffer phenol:chloroform:isoamyl alcohol protocol [2] and subject to isopycnic centrifugation in CsCl gradients. Briefly, 8 ml polyallomer tubes were filled with CsCl dissolved in Tris EDTA buffer (buoyant density 1.71 g $ml^{-1}$) and 6 µg genomic DNA before sealing and centrifugation at 152,000 x *g* in an MLN80 rotor (Beckman-Coulter) at 20°C for 72 h. CsCl gradients were then fractionated into ~350 µl aliquots before determining buoyant density via refractive index and recovery of DNA by PEG precipitation [2]. The predicted GC mol% of DNA in specific fractions was determined using the calculation of Schildkraut *et al.* [3].

*Quantitative PCR and metagenomic sequencing of fractionated DNA samples*

Quantitative PCR (qPCR) was performed to determine the distribution of prokaryotic and nitrifier genomes through CsCl gradients using a Corbett Rotor-Gene 6000 realtime PCR cycler. Prokaryotic 16S rRNA, bacterial and archaeal *amoA* genes were quantified across the entire buoyant density gradients using primer pairs P1(341f)/P2(534r) [4], amoA1F/amoA2R [5], and crenamoA23F/crenamoA616R [6]), respectively. Each 25 µl PCR contained 12.5 µl 2X QuantiFast SYBR Green Mix (Qiagen), 1 µM of each primer, 2 µl of standard DNA, recovered DNA from each fraction (1/10 dilution) or water (no template negative control). Thermal cycling programs consisted of an initial denaturation step of 15 min at 95°C, followed by 30 cycles of 15 s at 94°C, 30 s at 60°C, 30 s at 72°C for the 16S rRNA gene assay or 15 s at 94°C, 30 s at 55°C, 30 s at 72°C for bacterial and archaeal *amoA* genes assays followed by melt-curve analysis. All assays had an efficiency between 96 and 99% with an r2 value ≥0.99. After assessing the distribution of prokaryotic and nitrifier genomes, DNA in fractions with a buoyant density between 1.676 to 1.699 g $ml^{-1}$ (low buoyant density (LBD)) and 1.719 to 1.750 g $ml^{-1}$ (high buoyant density (HBD)) were pooled for metagenome sequencing. LBD and HBD metagenomes were sequenced at IntegraGen (Paris, France) and the Joint Genome Institute (JGI, Berkeley, CA, USA), respectively, both using the NovaSeq sequencing platform (Illumina) with NovaSeq XP version 1 reagent kits and a S4 flowcell 150 bp paired reads.

*Assembly, binning, and annotation*

Individual reads of HBD metagenomes were quality-trimmed with the MetaWrap read_qc module [7]. Co-assembly of the 135 to 245 million quality-controlled reads per metagenome was performed using MEGAHIT version 1.2.9 [8]. Binning was performed using metaBAT 2 [9], MaxBin2 [10] and CONCOCT [11] implemented in MetaWRAP version 1.2.1 [7] and bin completion and contamination estimated using CheckM version 1.0.12 [12]. The relative abundance of sequences mapping to MAG contigs in individual metagenomes was determined using default parameters in the MetaWRAP-Quant-bins module [7]. Taxonomic annotation of contigs was performed using Kaiju [13] with the NCBI nr database (2021-02-24). Taxonomy of MAGs was assigned using GTDB-Tk version 0.3.2 [14] with the Genome Taxonomy Database (GTDB, release date 2022-03-23).

*Identification of putative viruses infecting nitrifiers*

Viral contigs were predicted from contigs ≥10 kb using VirSorter [15], VirSorter 2.0 [16] and DeepVirFinder [17] (see Supplementary Figure 5 for comparison of predictions). CheckV [18] and manual curation (i.e. identification of viral hallmark genes such as capsid, portal, terminase and integrase proteins, enrichment in non-annotated genes) were performed to confirm a potential viral origin of each contig. The relative abundance of sequences mapping to virus contigs in individual metagenomes was determined using BBMap [19] with a threshold of ≥75% of contig length with ≥1x read coverage recruited at ≥90% average nucleotide identity [20]. The relative abundance of each viral contig in each HBD and LBD metagenome was calculated based on the length of sequence size and coverage and abundance was expressed as normalized copies per million reads (CPM). Gene prediction of each viral contig was performed using Prodigal version 2.6.3 with meta option [21], and homology search was performed against the NCBI nr database using Diamond blastp [22] and Interproscan5 [23].

To identify contigs from viruses infecting nitrifiers specifically, a database of virus genes present in proviruses of GTDB nitrifier reference genomes was constructed using VIBRANT [24], PhageBoost [25] and VirSorter [15] with 184 (87%) of 212 AOA genomes containing provirus regions. Capsid, terminase, portal and integrase protein sequences were then used as search queries against each viral contig using Diamond blastp (e-value < $10^{-5}$). Genes representing potential nitrifier-specific auxiliary metabolic genes were manually checked in annotation lists. A homologue-based approach was also used where an AOA host was predicted when 'best hit' shared homologues with those in genomes of *Thaumarchaeota* (NCBI taxonomy) were ≥3x more abundant compared to the second most dominant phylum [26]. Unlike our previous DNA-SIP analysis of methanotroph populations and associated viruses in these soils [27], spacer sequences in AOA MAG CRISPR arrays only matched predicted AOA viruses from other studies with a minimum of 2 mismatches between spacer and protospacer sequences and none from this study (data not shown).

*Phylogenomic, phylogenetic and predicted protein analyses*

Phylogenomic analysis of AOA MAGs and reference genomes from cultivated AOA and MAGs was performed using an alignment of single copy marker genes generated with GToTree [28] with a subsequent maximum likelihood phylogeny calculated using FastTree [29]. Single gene phylogenetic analyses were performed using alignments generated using MUSCLE [30] and manually refined. Maximum likelihood phylogenetic trees were constructed using PhyML [31] with automatic model selection. Affiliation of AOA MAGs to *amoA* gene-defined lineages was determined from phylogenetic analysis using the curated reference database of Alves *et al.* [32]. Where *amoA* genes were absent in incomplete *Nitrosotalea* MAGs, a broad-level affiliation was inferred from close phylogenomic relationships to complete genomes and their *amoA* lineage designation. Comparison of genome-wide similarity between AOA virus contigs and curated virus reference sequences was performed using ViPTree [33] and the Virus-Host DB [34] to generate a protein tree based on normalised tBLASTx scores. Virus gene-sharing network analyses were performed using vConTACT v2.0 [35].

**Supplementary Table 1.** Summary of metagenome sequencing of genomic DNA from low buoyant density (LBD) and high buoyant density (HBD) CsCl fractions derived from pH 4.5 and 7.5 soil. **A** Library accession numbers and read depth. **B** Contig assembly details.

**A**

| Sample ID* | NCBI BioProject Accession | Post-QC number of reads |
|---|---|---|
| LBD-12C-pH45_1 | PRJNA868779 | 170,519,430 |
| LBD-12C-pH45_2 | PRJNA868779 | 190,442,846 |
| LBD-12C-pH45_3 | PRJNA868779 | 183,142,278 |
| LBD-12C-pH75_1 | PRJNA868779 | 178,892,472 |
| LBD-12C-pH75_2 | PRJNA868779 | 180,678,072 |
| LBD-12C-pH75_3 | PRJNA868779 | 165,641,626 |
| HBD-12C-pH45_1 | PRJNA621418 | 137,927,826 |
| HBD-12C-pH45_2 | PRJNA621419 | 165,499,900 |
| HBD-12C-pH45_3 | PRJNA621420 | 161,786,148 |
| HBD-13C-pH45_1 | PRJNA621421 | 157,372,294 |
| HBD-13C-pH45_2 | PRJNA621422 | 140,157,446 |
| HBD-13C-pH45_3 | PRJNA621423 | 153,263,184 |
| HBD-12C-pH75_1 | PRJNA621424 | 172,033,988 |
| HBD-12C-pH75_2 | PRJNA621425 | 184,944,656 |
| HBD-12C-pH75_3 | PRJNA621426 | 173,060,618 |
| HBD-13C-pH75_1 | PRJNA621427 | 135,351,318 |
| HBD-13C-pH75_2 | PRJNA621428 | 245,188,756 |
| HBD-13C-pH75_3 | PRJNA621429 | 156,686,154 |

*name describes HBD or LBD CsCl fractions, $^{12}C$ or $^{13}C$ incubation, soil pH and replicate number.

**B**

| Sample ID | Assembled contigs | Mean contig length (bp) | Max. contig length (bp) |
|---|---|---|---|
| LBD-co-assembled contig | 26,611,526 | 614.2 | 809,772 |
| ≥5kb-LBD-co-assembled contig | 117,750 | 9,437.5 | |
| HBD-co-assembled contig | 35,109,680 | 566.3 | 812,893 |
| ≥5kb-HBD-co-assembled contig | 76,311 | 8,810.5 | |

**Supplementary Table 2.** Summary of 123 medium- and high-quality metagenome assembled genomes from low buoyant density DNA ranked by phylum (GTDB taxonomy).

| MAG ID | Completeness (%) | Contamination (%) | GC mol% | N50 (bp) | Size (bp) | Phylum | Class | Order | Family | Genus |
|---|---|---|---|---|---|---|---|---|---|---|
| 75 | 83.7 | 0.2 | 66.9 | 24,140 | 2,864,512 | Acidobacteriota | Acidobacteriae | Acidobacteriales | | |
| 56 | 82.5 | 0.0 | 60.3 | 15,391 | 3,027,569 | Acidobacteriota | Acidobacteriae | Acidobacteriales | Acidobacteriaceae | PALSA-350 |
| 101 | 80.7 | 5.1 | 54.0 | 21,626 | 3,847,244 | Acidobacteriota | Acidobacteriae | Acidobacteriales | Koribacteraceae | Gp1-AA145 |
| 126 | 77.1 | 3.0 | 67.5 | 17,321 | 3,322,481 | Acidobacteriota | Vicinamibacteria | Vicinamibacterales | UBA2999 | |
| 72 | 73.9 | 0.9 | 58.4 | 14,749 | 2,220,118 | Acidobacteriota | Acidobacteriae | UBA7541 | UBA7541 | |
| 124 | 67.3 | 3.4 | 63.9 | 11,528 | 2,512,459 | Acidobacteriota | Acidobacteriae | Acidobacteriales | | |
| 33 | 66.0 | 5.5 | 59.3 | 15,405 | 2,887,037 | Acidobacteriota | Acidobacteriae | UBA7541 | UBA7541 | Palsa-295 |
| 22 | 53.3 | 0.9 | 56.3 | 9,893 | 2,452,695 | Acidobacteriota | Acidobacteriae | UBA7541 | UBA7541 | |
| 68 | 50.8 | 3.4 | 61.9 | 9,900 | 2,500,689 | Acidobacteriota | Acidobacteriae | Acidobacteriales | Acidobacteriaceae | KBS-83 |
| 87 | 50.4 | 2.0 | 52.4 | 8,779 | 1,812,072 | Acidobacteriota | Blastocatellia | Pyrinomonadales | Pyrinomonadaceae | OLB17 |
| 111 | 97.3 | 1.9 | 66.6 | 36,117 | 2,871,784 | Actinobacteriota | Thermoleophilia | Solirubrobacterales | Solirubrobacteraceae | |
| 44 | 97.0 | 2.6 | 66.0 | 142,875 | 3,274,599 | Actinobacteriota | Thermoleophilia | Solirubrobacterales | Solirubrobacteraceae | |
| 25 | 97.0 | 3.1 | 65.7 | 71,608 | 2,181,555 | Actinobacteriota | Thermoleophilia | Solirubrobacterales | 70-9 | |
| 46 | 82.0 | 0.1 | 63.9 | 17,436 | 1,359,101 | Actinobacteriota | Acidimicrobiia | Acidimicrobiales | RAAP-2 | RAAP-2 |
| 6 | 79.2 | 2.2 | 68.5 | 54,707 | 2,098,665 | Actinobacteriota | Thermoleophilia | Solirubrobacterales | 70-9 | |
| 57 | 70.8 | 1.2 | 69.6 | 12,884 | 2,594,437 | Actinobacteriota | Actinobacteria | Nanopelagicales | | |
| 48 | 63.0 | 3.0 | 70.1 | 11,121 | 2,066,628 | Actinobacteriota | Acidimicrobiia | Acidimicrobiales | RAAP-2 | Bog-756 |
| 93 | 60.3 | 0.9 | 62.8 | 10,343 | 1,801,421 | Actinobacteriota | Acidimicrobiia | Microtrichales | Ilumatobacteraceae | UBA668 |
| 79 | 67.5 | 0.9 | 61.7 | 11,258 | 2,006,440 | Armatimonadota | Fimbriimonadia | Fimbriimonadales | Fimbriimonadaceae | |
| 38 | 96.5 | 0.7 | 44.9 | 52,178 | 3,501,364 | Bacteroidota | Bacteroidia | Chitinophagales | Chitinophagaceae | JJ008 |
| 100 | 94.8 | 1.5 | 39.6 | 298,057 | 4,276,415 | Bacteroidota | Bacteroidia | AKYH767 | | |
| 53 | 92.0 | 1.7 | 44.7 | 26,881 | 3,591,767 | Bacteroidota | Bacteroidia | Chitinophagales | Chitinophagaceae | UBA8621 |
| 62 | 86.9 | 7.5 | 38.5 | 28,454 | 3,315,570 | Bacteroidota | Bacteroidia | AKYH767-A | OLB10 | |
| 76 | 84.0 | 0.7 | 49.6 | 26,115 | 3,041,112 | Bacteroidota | Bacteroidia | AKYH767 | 2-12-FULL-35-15 | |
| 97 | 78.6 | 2.1 | 39.6 | 15,282 | 3,408,942 | Bacteroidota | Bacteroidia | AKYH767 | | |
| 81 | 77.5 | 0.2 | 54.5 | 15,405 | 2,291,910 | Bacteroidota | Bacteroidia | Chitinophagales | Chitinophagaceae | |
| 106 | 76.6 | 3.4 | 44.4 | 12,939 | 2,749,566 | Bacteroidota | Bacteroidia | AKYH767 | b-17BO | PALSA-968 |
| 32 | 75.2 | 3.8 | 40.6 | 13,274 | 4,299,582 | Bacteroidota | Bacteroidia | AKYH767-A | OLB10 | |
| 47 | 73.6 | 4.2 | 34.8 | 11,826 | 2,228,512 | Bacteroidota | Ignavibacteria | Ignavibacteriales | Ignavibacteriaceae | BMS3ABIN03 |
| 116 | 73.2 | 1.9 | 51.5 | 13,585 | 2,332,193 | Bacteroidota | Kapabacteria | Palsa-1295 | Palsa-1295 | |
| 13 | 71.5 | 0.0 | 42.1 | 13,719 | 2,460,739 | Bacteroidota | Bacteroidia | NS11-12g | UBA955 | |
| 30 | 67.2 | 0.5 | 37.7 | 10,543 | 1,634,027 | Bacteroidota | Bacteroidia | Sphingobacteriales | Sphingobacteriaceae | |
| 26 | 66.5 | 1.5 | 42.0 | 11,489 | 3,546,402 | Bacteroidota | Bacteroidia | Chitinophagales | Chitinophagaceae | |
| 86 | 65.7 | 3.4 | 34.7 | 9,330 | 2,102,202 | Bacteroidota | Bacteroidia | Flavobacteriales | Flavobacteriaceae | Flavobacterium |
| 89 | 65.4 | 0.0 | 39.1 | 18,038 | 2,305,465 | Bacteroidota | Ignavibacteria | SJA-28 | OLB5 | |
| 40 | 64.5 | 7.3 | 39.0 | 8,876 | 2,679,712 | Bacteroidota | Bacteroidia | Chitinophagales | Chitinophagaceae | Taibaiella_B |
| 118 | 52.0 | 3.2 | 38.0 | 9,613 | 3,001,274 | Bacteroidota | Bacteroidia | Chitinophagales | Chitinophagaceae | Palsa-955 |
| 41 | 51.3 | 4.4 | 39.4 | 8,197 | 2,857,311 | Bacteroidota | Bacteroidia | Chitinophagales | BACL12 | UBA7236 |
| 83 | 50.8 | 1.0 | 41.9 | 11,602 | 2,718,158 | Bacteroidota | Bacteroidia | Chitinophagales | Chitinophagaceae | |
| 49 | 52.6 | 1.8 | 42.5 | 10,353 | 1,916,736 | Bdellovibrionota | Bdellovibrionia | Bdellovibrionales | Bdellovibrionaceae | Bdellovibrio |
| 58 | 91.4 | 0.0 | 61.2 | 14,550 | 3,678,188 | Chloroflexota | UBA5177 | UBA5177 | UBA5177 | |
| 69 | 75.9 | 2.2 | 62.3 | 12,810 | 2,934,925 | Chloroflexota | UBA4733 | UBA4733 | UBA4733 | |
| 115 | 75.1 | 2.8 | 70.3 | 18,220 | 2,481,858 | Chloroflexota | Ellin6529 | CSP1-4 | CSP1-4 | Palsa-1033 |
| 16 | 64.2 | 1.4 | 55.1 | 23,141 | 1,640,525 | Chloroflexota | Anaerolineae | Anaerolineales | UBA11579 | |
| 121 | 63.5 | 8.2 | 48.7 | 8,645 | 3,693,765 | Chloroflexota | Anaerolineae | Anaerolineales | envOPS12 | OLB14 |
| 14 | 62.9 | 1.0 | 68.7 | 11,300 | 3,380,230 | Chloroflexota | Ktedonobacteria | Ktedonobacterales | | |

| | | | | | | | | | | |
|---|---|---|---|---|---|---|---|---|---|---|
| 36 | 59.5 | 2.8 | 72.0 | 10,127 | 1,826,549 | Chloroflexota | Ellin6529 | CSP1-4 | CSP1-4 | CSP1-4 |
| 17 | 57.0 | 0.9 | 64.1 | 9,099 | 3,136,804 | Chloroflexota | UBA5177 | | | |
| 24 | 56.3 | 0.9 | 63.3 | 7,671 | 2,328,854 | Chloroflexota | UBA5177 | | | |
| 43 | 92.3 | 0.9 | 43.4 | 23,016 | 2,895,603 | Cyanobacteria | Vampirovibrionia | Vampirovibrionales | | |
| 104 | 52.6 | 0.0 | 46.6 | 9,501 | 1,686,795 | Cyanobacteria | Vampirovibrionia | Vampirovibrionales | | |
| 84 | 62.1 | 0.0 | 35.2 | 9,392 | 965,279 | Dependentiae | Babeliae | Babeliales | | |
| 11 | 60.2 | 0.9 | 69.4 | 14,745 | 1,971,573 | Dormibacterota | Dormibacteria | UBA8260 | UBA8260 | |
| 23 | 54.3 | 9.5 | 68.9 | 8,592 | 1,357,015 | Dormibacterota | Dormibacteria | UBA8260 | UBA8260 | Palsa-851 |
| 119 | 87.2 | 0.0 | 38.0 | 15,691 | 2,341,565 | Eremiobacterota | Eremiobacteria | Eremiobacterales | Eremiobacteraceae | |
| 103 | 77.4 | 1.5 | 68.2 | 12,265 | 2,115,044 | Eremiobacterota | Eremiobacteria | UBP12 | UBA5184 | |
| 37 | 51.1 | 7.4 | 60.2 | 7,077 | 1,974,572 | Eremiobacterota | Eremiobacteria | UBP12 | UBA5184 | PALSA-1484 |
| 82 | 95.1 | 0.3 | 34.9 | 35,039 | 2,302,984 | Firmicutes | Clostridia | Lachnospirales | Lachnospiraceae | Herbinix |
| 70 | 84.4 | 2.6 | 41.4 | 17,334 | 3,226,646 | Firmicutes | Bacilli | Bacillales | Bacillaceae | |
| 10 | 78.7 | 0.8 | 45.6 | 21,297 | 2,470,344 | Firmicutes | Bacilli | Bacillales | Bacillaceae | |
| 123 | 59.2 | 1.0 | 36.4 | 18,393 | 979,880 | Firmicutes | Bacilli | Bacillales | Planococcaceae | Ureibacillus |
| 99 | 54.5 | 9.9 | 36.8 | 6,988 | 796,782 | Firmicutes | Bacilli | Bacillales | Bacillaceae_G | Bacillus |
| 3 | 51.9 | 5.3 | 32.6 | 12,001 | 1,498,041 | Firmicutes | Clostridia | Acetivibrionales | Acetivibrionaceae | Herbivorax |
| 80 | 97.4 | 0.7 | 41.7 | 35,221 | 2,034,588 | Firmicutes_F | Halanaerobiia | Halanaerobiales | DTU029 | DTU029 |
| 35 | 99.2 | 1.0 | 42.0 | 79,014 | 2,517,665 | Firmicutes_I | Bacilli | Thermoactinomycetales | Thermoactinomycetaceae | |
| 120 | 96.5 | 0.3 | 48.2 | 35,048 | 2,270,355 | Firmicutes_I | Bacilli | Thermoactinomycetales | Thermoactinomycetaceae | CDF |
| 67 | 69.4 | 0.3 | 41.6 | 9,016 | 1,357,832 | Firmicutes_I | Bacilli | Thermoactinomycetales | Thermoactinomycetaceae | |
| 122 | 71.8 | 2.7 | 65.3 | 16,326 | 2,944,945 | Gemmatimonadota | Gemmatimonadetes | Gemmatimonadales | Gemmatimonadaceae | FEN-1250 |
| 34 | 71.6 | 3.4 | 66.1 | 29,597 | 3,506,635 | Gemmatimonadota | Gemmatimonadetes | Gemmatimonadales | Gemmatimonadaceae | |
| 39 | 69.1 | 1.2 | 67.2 | 17,110 | 3,202,677 | Gemmatimonadota | Gemmatimonadetes | Gemmatimonadales | Gemmatimonadaceae | |
| 91 | 52.7 | 0.0 | 62.8 | 9,236 | 2,678,084 | Gemmatimonadota | Gemmatimonadetes | Gemmatimonadales | Gemmatimonadaceae | AG2 |
| 117 | 58.9 | 0.0 | 68.6 | 7,421 | 2,581,217 | Myxococcota | Polyangia | Palsa-1104 | Palsa-1104 | PALSA-1104 |
| 31 | 78.3 | 0.0 | 35.0 | 63,246 | 1,110,445 | Patescibacteria | Dojkabacteria | | | |
| 110 | 72.5 | 0.0 | 45.6 | 538,338 | 833,363 | Patescibacteria | Saccharimonadia | Saccharimonadales | 2-12-FULL-41-12 | |
| 102 | 70.0 | 0.0 | 35.8 | 24,450 | 1,170,104 | Patescibacteria | Microgenomatia | Levybacterales | UBA12049 | |
| 60 | 69.9 | 1.9 | 63.1 | 809,772 | 809,772 | Patescibacteria | Saccharimonadia | | | |
| 51 | 65.0 | 0.8 | 58.3 | 36,742 | 983,527 | Patescibacteria | Paceibacteria | UBA6257 | 2-01-FULL-56-20 | |
| 50 | 64.6 | 0.0 | 39.7 | 277,019 | 821,951 | Patescibacteria | Microgenomatia | UBA1406 | HO2-37-13b | |
| 66 | 64.5 | 0.0 | 41.9 | 35,340 | 846,808 | Patescibacteria | Microgenomatia | UBA1406 | GWC2-37-13 | 2-01-FULL-40-42 |
| 42 | 64.4 | 0.0 | 43.7 | 72,802 | 632,826 | Patescibacteria | Saccharimonadia | Saccharimonadales | UBA10212 | |
| 7 | 64.2 | 0.0 | 53.8 | 40,147 | 643,112 | Patescibacteria | Saccharimonadia | Saccharimonadales | UBA10212 | |
| 29 | 60.9 | 1.0 | 43.9 | 26,687 | 688,051 | Patescibacteria | Doudnabacteria | UBA920 | O2-02-FULL-48-8 | |
| 45 | 59.9 | 0.0 | 39.8 | 12,821 | 852,036 | Patescibacteria | Microgenomatia | Levybacterales | UBA12049 | PRDT01 |
| 28 | 59.4 | 0.0 | 49.9 | 16,270 | 829,123 | Patescibacteria | Saccharimonadia | Saccharimonadales | UBA4665 | UBA6224 |
| 74 | 54.6 | 0.0 | 35.5 | 26,797 | 536,454 | Patescibacteria | Microgenomatia | Levybacterales | UBA12049 | |
| 114 | 53.4 | 6.5 | 43.9 | 9,739 | 841,384 | Patescibacteria | Saccharimonadia | Saccharimonadales | 2-12-FULL-41-12 | |
| 27 | 52.7 | 0.4 | 46.8 | 40,409 | 442,116 | Patescibacteria | Paceibacteria | UBA9983 | Zambryskibacteraceae | UBA5004 |
| 5 | 51.9 | 0.0 | 37.5 | 13,915 | 425,834 | Patescibacteria | Paceibacteria | UBA9983 | Zambryskibacteraceae | C7867-006 |
| 54 | 50.4 | 0.0 | 38.2 | 21,102 | 760,548 | Patescibacteria | Microgenomatia | Woykebacterales | | |
| 8 | 66.5 | 0.0 | 58.4 | 10,752 | 3,564,280 | Planctomycetota | Phycisphaerae | UBA1161 | UBA1161 | |
| 105 | 65.4 | 5.8 | 59.4 | 11,513 | 4,165,716 | Planctomycetota | Phycisphaerae | UBA1161 | UBA1161 | |
| 73 | 95.1 | 4.3 | 65.6 | 137,474 | 4,398,083 | Proteobacteria | Gammaproteobacteria | Steroidobacterales | Steroidobacteraceae | |
| 19 | 89.5 | 2.6 | 70.1 | 31,514 | 3,812,969 | Proteobacteria | Alphaproteobacteria | Caulobacterales | Caulobacteraceae | BOG-938 |
| 125 | 88.0 | 3.1 | 62.4 | 28,686 | 2,191,964 | Proteobacteria | Alphaproteobacteria | Sphingomonadales | Sphingomonadaceae | Sphingomonas |
| 108 | 87.5 | 6.2 | 57.0 | 22,826 | 1,972,878 | Proteobacteria | Alphaproteobacteria | Micavibrionales | Micavibrionaceae | UM-FILTER-47-13 |
| 92 | 82.8 | 5.0 | 65.9 | 19,149 | 2,660,651 | Proteobacteria | Gammaproteobacteria | Steroidobacterales | Steroidobacteraceae | 13-2-20CM-66-19 |
| 64 | 81.0 | 1.7 | 61.6 | 22,864 | 2,553,403 | Proteobacteria | Gammaproteobacteria | Xanthomonadales | Rhodanobacteraceae | Rudaea |
| 71 | 79.5 | 0.9 | 61.2 | 22,385 | 2,981,098 | Proteobacteria | Alphaproteobacteria | UBA1301 | UBA1301 | UBA6038 |

| | | | | | | | | | | |
|---|---|---|---|---|---|---|---|---|---|---|
| 98 | 79.3 | 2.1 | 70.5 | 22,416 | 2,489,191 | Proteobacteria | Gammaproteobacteria | Steroidobacterales | Steroidobacteraceae | 13-2-20CM-66-19 |
| 94 | 78.7 | 0.9 | 59.3 | 14,277 | 2,998,862 | Proteobacteria | Alphaproteobacteria | Rhizobiales | Xanthobacteraceae | Pseudolabrys |
| 95 | 77.7 | 0.0 | 35.2 | 13,606 | 3,485,685 | Proteobacteria | Gammaproteobacteria | UBA5158 | UBA5158 | |
| 65 | 77.3 | 1.1 | 66.2 | 16,642 | 5,023,705 | Proteobacteria | Alphaproteobacteria | Acetobacterales | Acetobacteraceae | Palsa-883 |
| 2 | 73.3 | 1.4 | 67.0 | 20,592 | 2,309,631 | Proteobacteria | Alphaproteobacteria | Sphingomonadales | Sphingomonadaceae | Porphyrobacter |
| 109 | 61.9 | 5.3 | 63.3 | 13,874 | 3,496,047 | Proteobacteria | Alphaproteobacteria | Rhizobiales | Xanthobacteraceae | BOG-931 |
| 78 | 56.7 | 1.3 | 64.9 | 35,205 | 1,728,632 | Proteobacteria | Alphaproteobacteria | Elsterales | URHD0088 | |
| 55 | 56.4 | 0.8 | 61.4 | 11,298 | 1,368,874 | Proteobacteria | Alphaproteobacteria | Rhizobiales | Methyloligellaceae | Methyloceanibacter |
| 90 | 55.5 | 2.1 | 67.2 | 11,614 | 1,755,960 | Proteobacteria | Gammaproteobacteria | Burkholderiales | Palsa-1005 | PALSA-1003 |
| 88 | 55.2 | 3.4 | 65.1 | 9,131 | 1,518,294 | Proteobacteria | Alphaproteobacteria | Rhizobiales | Xanthobacteraceae | Palsa-892 |
| 18 | 52.5 | 5.7 | 59.0 | 7,355 | 2,371,465 | Proteobacteria | Alphaproteobacteria | Rhizobiales | Methyloligellaceae | Methyloceanibacter |
| 20 | 51.7 | 0.0 | 61.0 | 9,057 | 2,689,297 | Proteobacteria | Alphaproteobacteria | UBA1301 | UBA1301 | |
| 15 | 51.2 | 1.7 | 59.9 | 9,856 | 2,680,085 | Proteobacteria | Alphaproteobacteria | Rhizobiales | Xanthobacteraceae | Nitrobacter |
| 21 | 50.3 | 1.1 | 64.3 | 8,593 | 1,496,296 | Proteobacteria | Gammaproteobacteria | Burkholderiales | Burkholderiaceae | C04 |
| 96 | 91.3 | 1.9 | 31.0 | 10,635 | 4,039,103 | Thermoproteota | Nitrososphaeria | Nitrososphaerales | Nitrososphaeraceae | TH1177 |
| 52 | 88.8 | 1.9 | 49.7 | 15,865 | 1,740,480 | Thermoproteota | Nitrososphaeria | Nitrososphaerales | Nitrososphaeraceae | Nitrososphaera |
| 61 | 87.9 | 1.9 | 37.8 | 14,936 | 1,491,779 | Thermoproteota | Nitrososphaeria | Nitrososphaerales | Nitrosopumilaceae | Nitrosotalea |
| 107 | 85.9 | 2.9 | 37.8 | 36,213 | 1,658,330 | Thermoproteota | Nitrososphaeria | Nitrososphaerales | Nitrosopumilaceae | Nitrosotalea |
| 63 | 78.6 | 1.0 | 37.4 | 10,792 | 2,066,632 | Thermoproteota | Nitrososphaeria | Nitrososphaerales | Nitrososphaeraceae | |
| 12 | 73.3 | 3.2 | 50.2 | 11,601 | 1,019,284 | Thermoproteota | Nitrososphaeria | Nitrososphaerales | Nitrososphaeraceae | Nitrososphaera |
| 77 | 65.2 | 6.5 | 36.8 | 16,902 | 1,553,150 | Thermoproteota | Nitrososphaeria | Nitrososphaerales | Nitrosopumilaceae | Nitrosotalea |
| 112 | 62.9 | 1.0 | 37.9 | 9,638 | 1,212,031 | Thermoproteota | Nitrososphaeria | Nitrososphaerales | Nitrososphaeraceae | TA-21 |
| 9 | 61.7 | 1.0 | 37.3 | 13,015 | 970,273 | Thermoproteota | Nitrososphaeria | Nitrososphaerales | Nitrosopumilaceae | Nitrosotalea |
| 85 | 98.6 | 8.6 | 57.1 | 36,458 | 4,399,804 | Verrucomicrobiota | Verrucomicrobiae | Pedosphaerales | Pedosphaeraceae | UBA11358 |
| 59 | 94.6 | 1.6 | 37.8 | 26,369 | 2,935,834 | Verrucomicrobiota | Chlamydiia | Parachlamydiales | Parachlamydiaceae | |

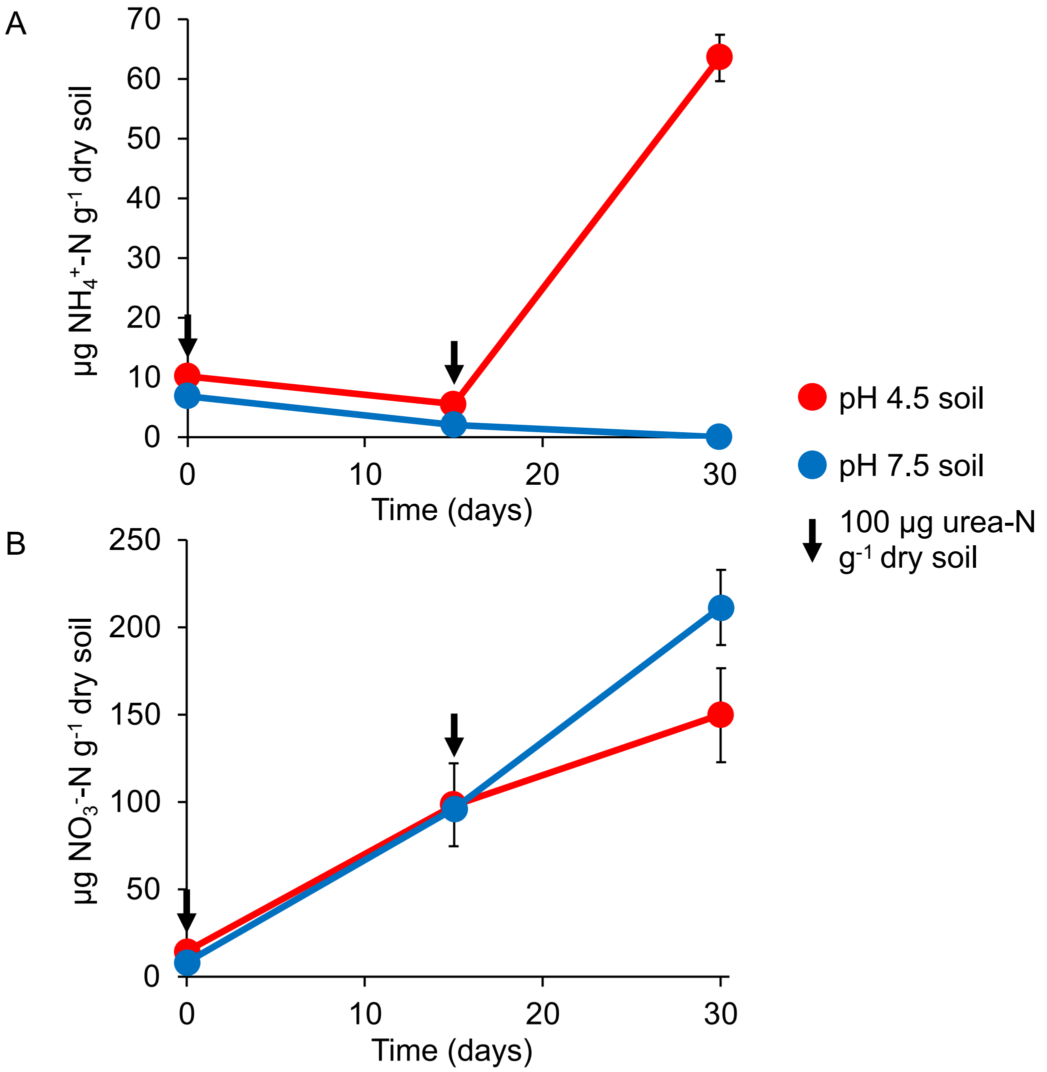


**Supplementary Figure 1.** Nitrification kinetics in soil microcosms. **A** Ammonium and **B** Nitrite+nitrate concentrations in pH 4.5 and 7.5 soil microcosms after addition of 2 x 100 µg urea-N $g^{-1}$ soil. Ammonia concentrations were determined prior to the addition of urea on day 0 and 15. Points and error bars represent the mean and standard error value from three destructively sampled microcosms, with some error bars smaller than the symbol. Microcosms were incubated with $^{12}C$-$CO_2$ for determining nitrification kinetics with no significant differences observed with N concentrations in $^{13}C$ microcosms at day 30 (data not shown).

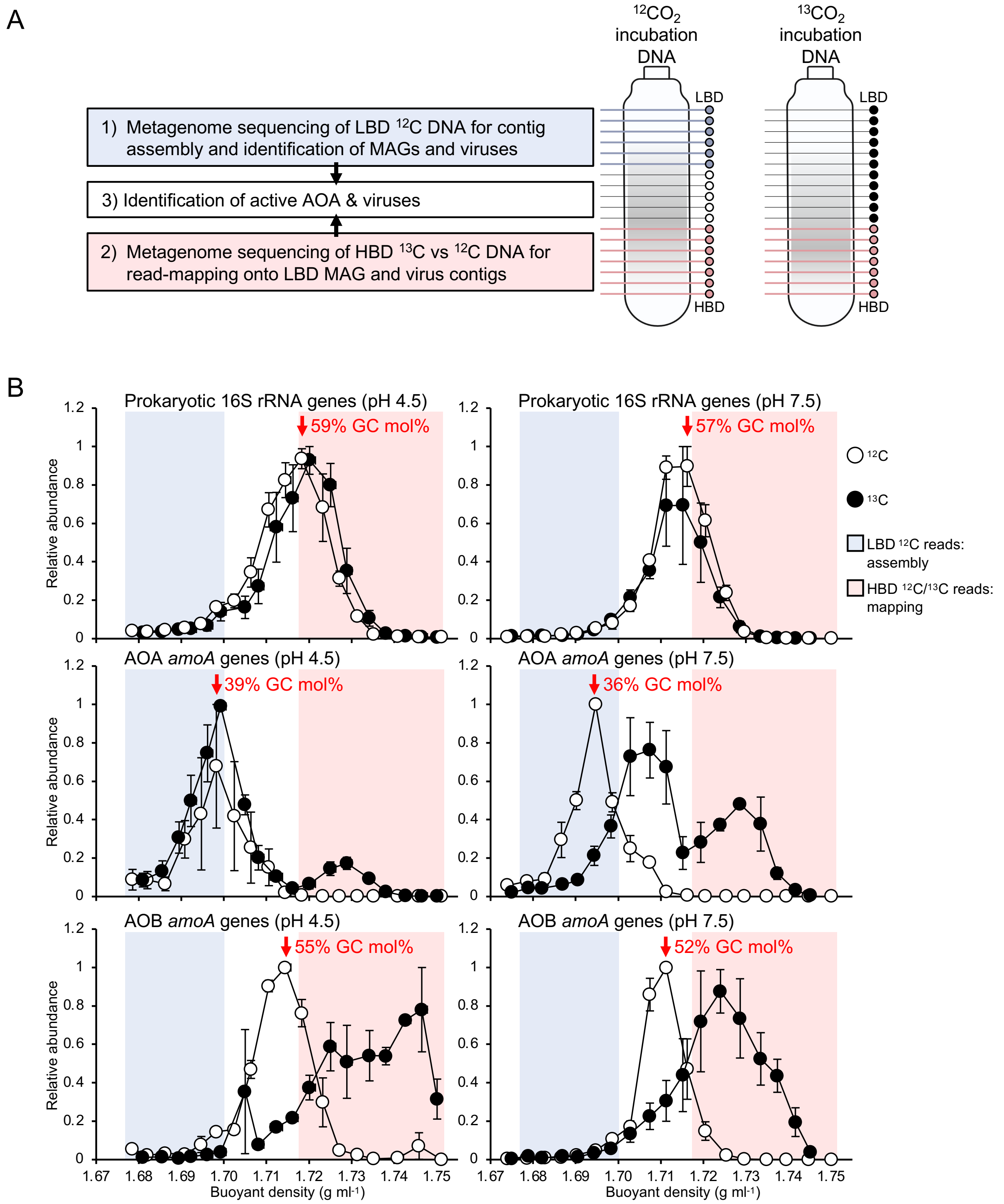


**Supplementary Figure 2.** Isopycnic centrifugation and metagenomic sequencing of DNA in targeted fractions for hybrid analysis of $^{12}C$- and $^{13}C$-enriched DNA. **A** Schematic of GC mol% fractionation in CsCl and selection of samples for metagenomic sequencing and analysis. **B** Distribution of total prokaryotic, AOA and AOB genomes in CsCl gradients determined from the relative abundance of 16S rRNA genes (prokaryotes) and *amoA* genes (AOA and AOB). Vertical error bars are the standard error of the mean relative abundance and horizontal bars (mostly smaller than the symbol size) the standard error of the mean buoyant density of individual fractions from three independent CsCl gradients, each representing an individual microcosm. Fractions highlighted in blue or pink areas were pooled for each replicate microcosm for sequencing. The GC mol% of genomic DNA is given (red text and arrow) for the $^{12}C$ fraction containing the highest quantity of genomic DNA for each target group.

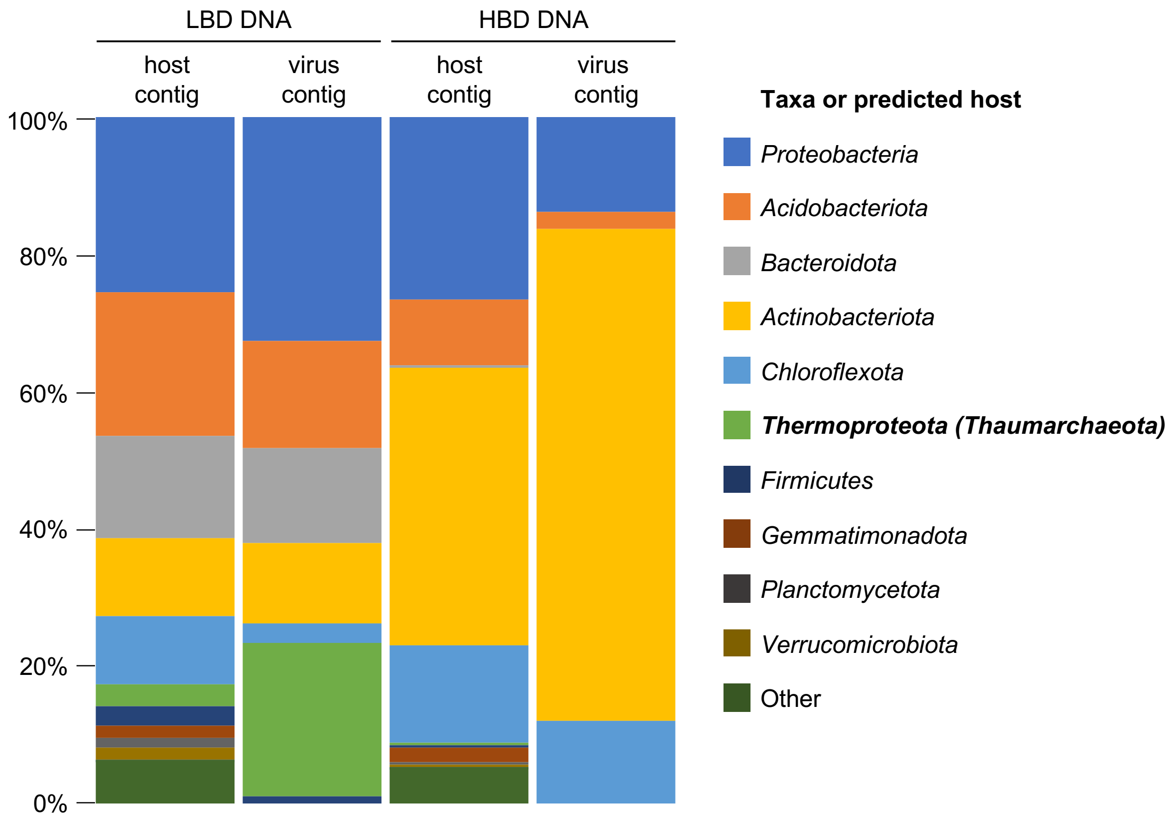


**Supplementary Figure 3.** Taxonomic affiliation (GTDB) of contigs from prokaryote genomes (host contigs) or predicted hosts of viruses (virus contigs) in low buoyant density (LBD) and high buoyant density (HBD) metagenomic libraries. Unaffiliated contigs are not included. AOA of the class *Nitrososphaeria* are placed within the *Thermoproteota* (GTDB), *Nitrososphaerota* (ICNP) or *Thaumarchaeota* (NCBI) phylum. Data are the mean values from both pH 4.5 and 7.5 libraries.

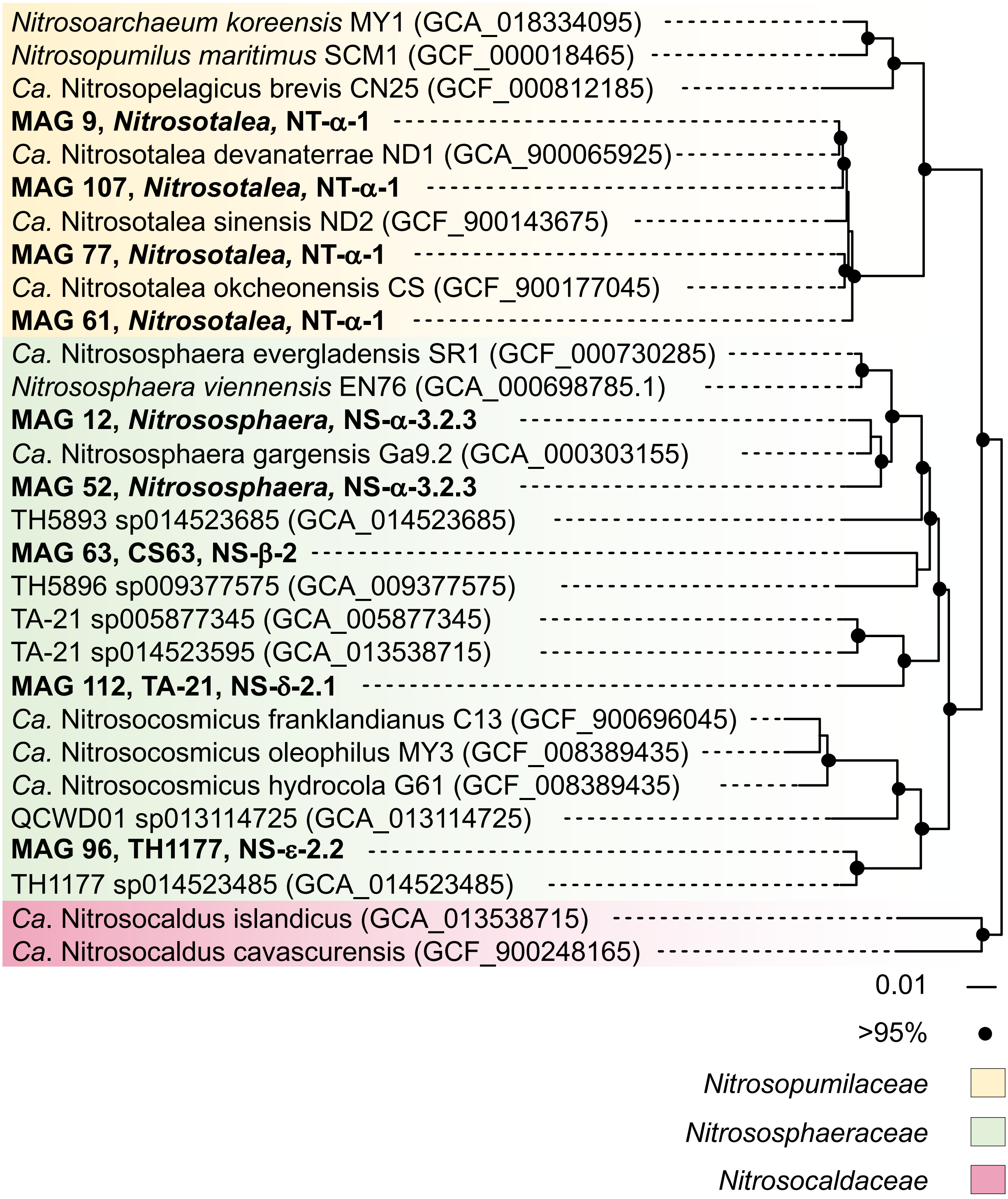


**Supplementary Figure 4.** Maximum likelihood phylogenomic analysis of nine AOA MAGs within the class *Nitrososphaeria*. MAGs were compared with cultivated AOA genomes or reference MAGs using 14,524 aligned positions from 76 single copy genes and rooted with *Nitrosocaldus* representatives. NCBI accession numbers are given in parenthesis. Circles at nodes represent >95% percentage bootstrap support from 1000 replicates and the scale bar denotes an estimated 0.01 changes per position.

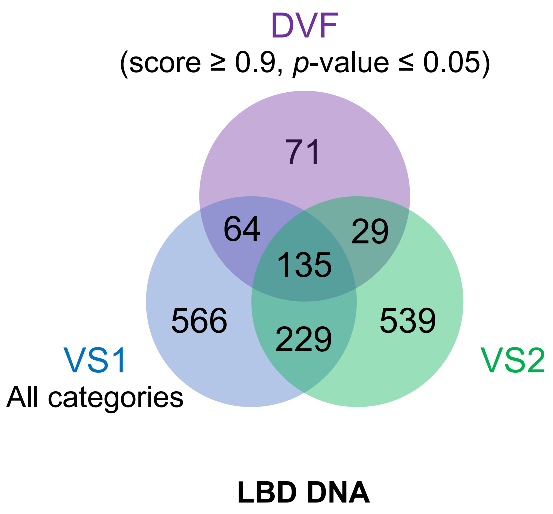


**Supplementary Figure 5.** Venn diagrams comparing the number of predicted virus contigs using VirSorter1, VirSorter2 and DeepVirFinder tools from low buoyant density (LBD) and high buoyant density (HBD) DNA metagenomic libraries from both pH 4.5 and 7.5 soils.
